## Supplementary Figures for "GATA2 regulates mast cell identity and responsiveness to antigenic stimulation by promoting chromatin remodeling at super-enhancers"

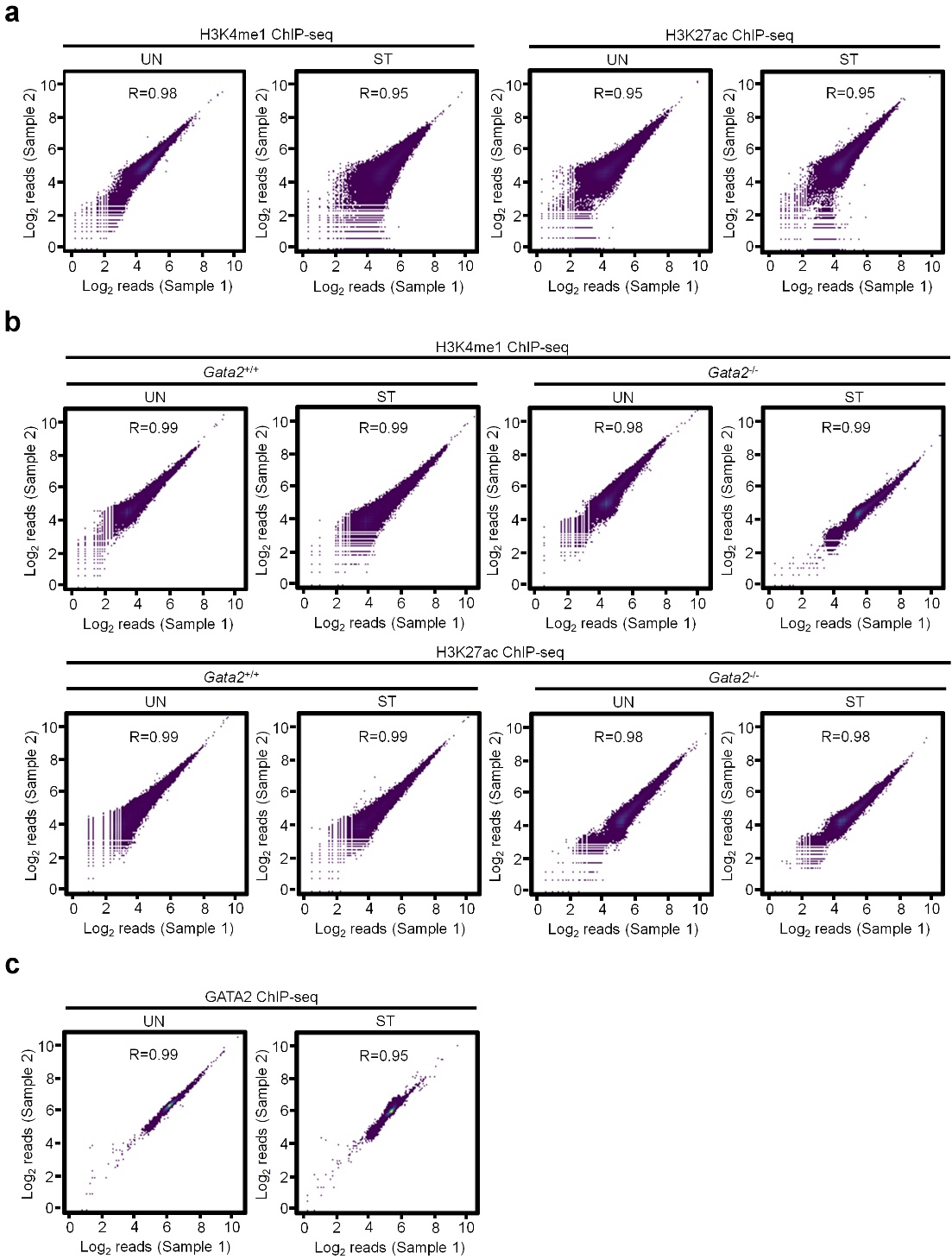

**Supplementary Figure 1 (related to Fig 1 to Fig 7). Reproducibility of the H3K4me1, H3K27ac and GATA2 ChIP-seq data. a** The reproducibility of H3K4me1 and H3K27ac ChIP-seq data from two biological replicates in resting (UN) and Stimulated (ST) BMMCs. **b** The reproducibility of H3K4me1 and H3K27ac ChIP-seq data from two biological replicates in resting (UN) and Stimulated (ST) *Gata2* KO (*Gata2*^-/-^) or WT (*Gata2*^+/+^) BMMCs. **c** The reproducibility of GATA2 ChIP-seq data from two biological replicates in resting (UN) and Stimulated (ST) BMMCs. Reproducibility of the ChIP-seq data was analyzed using the deepTools (3.3.0) multiBamSummary and plotCorrelation functions. R values were calculated using Pearson correlation coefficient.

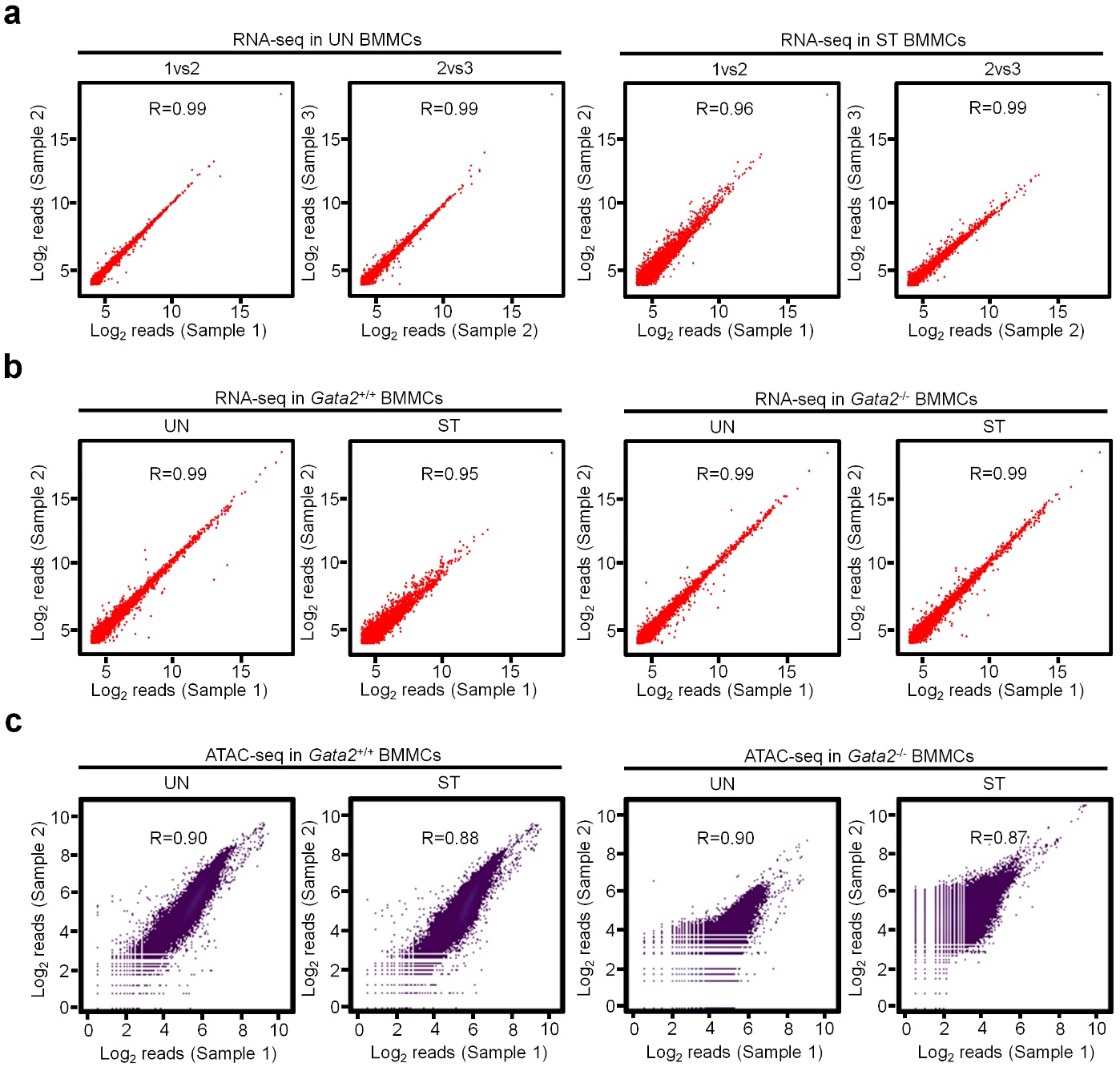

**Supplementary Figure 2 (related to Fig 1 to Fig 7). Reproducibility of the RNA-seq and Omni-ATAC-seq data. a** The reproducibility of RNA-seq data from three biological replicates in resting (UN) and Stimulated (ST) BMMCs. **b** The reproducibility of RNA-seq data from two biological replicates in resting (UN) and Stimulated (ST) *Gata2* KO (*Gata2*^-/-^) or WT (*Gata2*^+/+^) BMMCs. **c** The reproducibility of Omni-ATAC-seq data from two biological replicates in resting (UN) and Stimulated (ST) *Gata2* KO (*Gata2*^-/-^) or WT (*Gata2*^+/+^) BMMCs. Reproducibility of the RNA-seq data was calculated using the ggscatter function in R package ggpubr (0.2). Reproducibility of the Moni-ATAC-seq data was analyzed using the deepTools (3.3.0) multiBamSummary and plotCorrelation functions. R values were calculated using Pearson correlation coefficient.

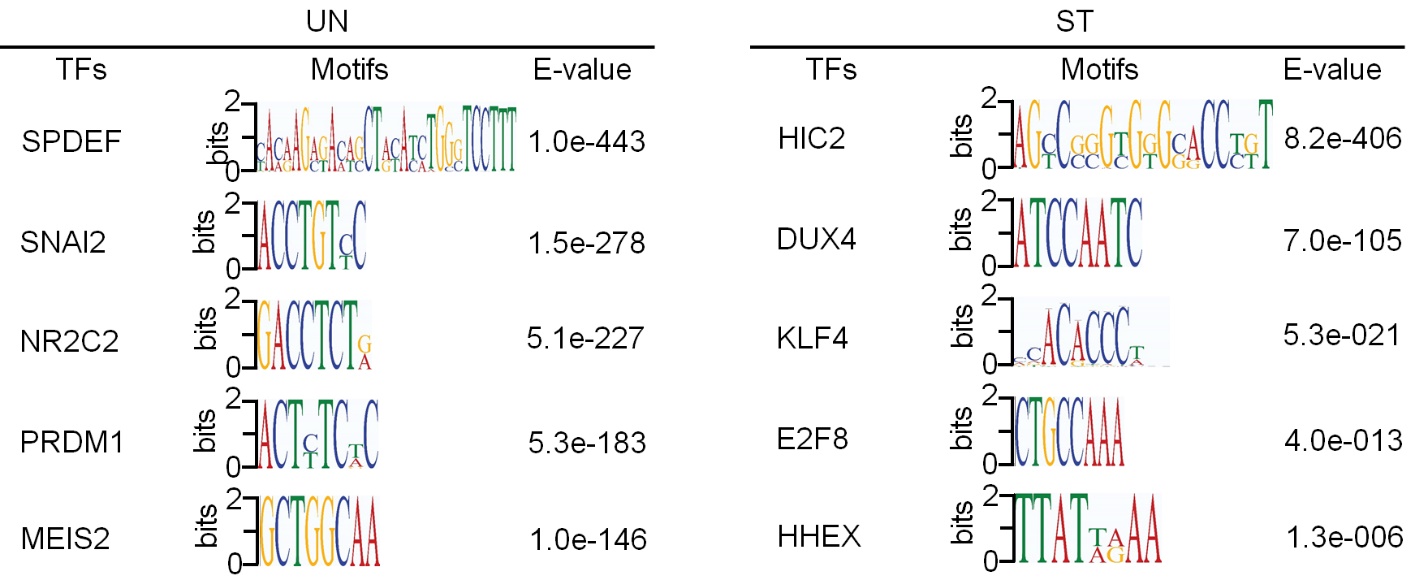

**Supplementary Figure 3 (related to Fig 3 and Fig 7). TF binding motif enrichment analysis.** TF binding motif enrichment analysis performed on the accessible regions located outside of the regions with H3K27ac modification in resting mast cells (UN, left panel) and the accessible regions located outside of the regions with increase H3K27ac modification in stimulated mast cells (ST, right panel).

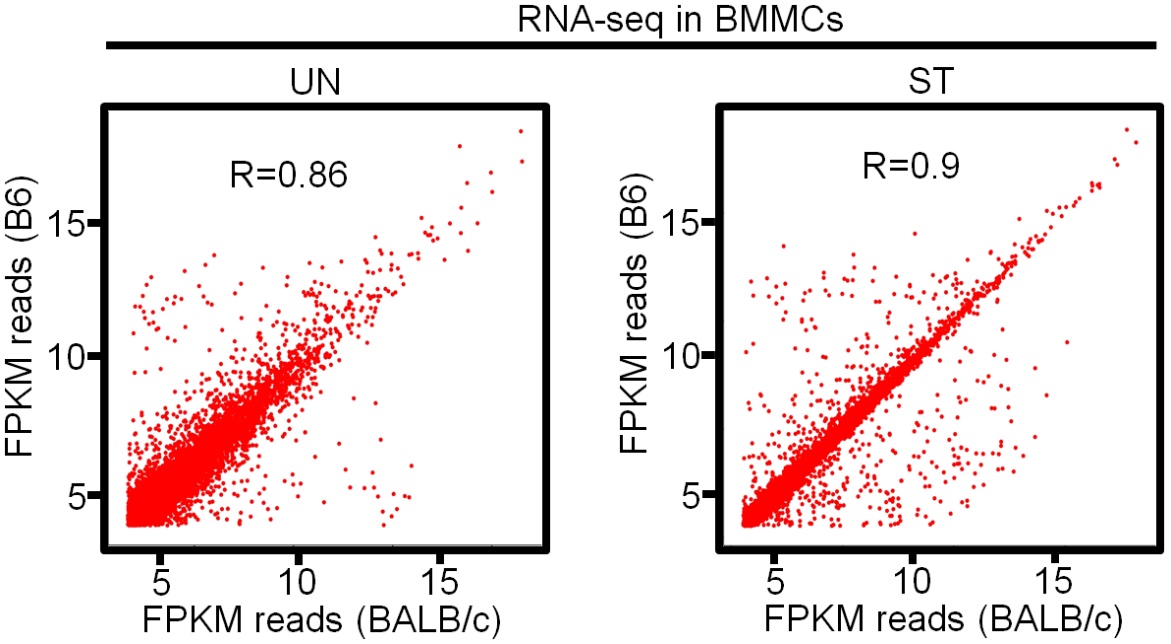

**Supplementary Figure 4 (related to Fig 3). Similarity of the gene expression profiles in C57BL/6 (B6) BMMCs and BALB/c BMMCs in resting (UN) and stimulated (ST) conditions.** The log2 ratio of FPKM reads of genes in BALB/c versus B6 BMMCs under resting (UN) or stimulated (ST) conditions were plotted by the ggscatter function in R package ggpubr (0.2). R values were calculated using Pearson correlation coefficient.

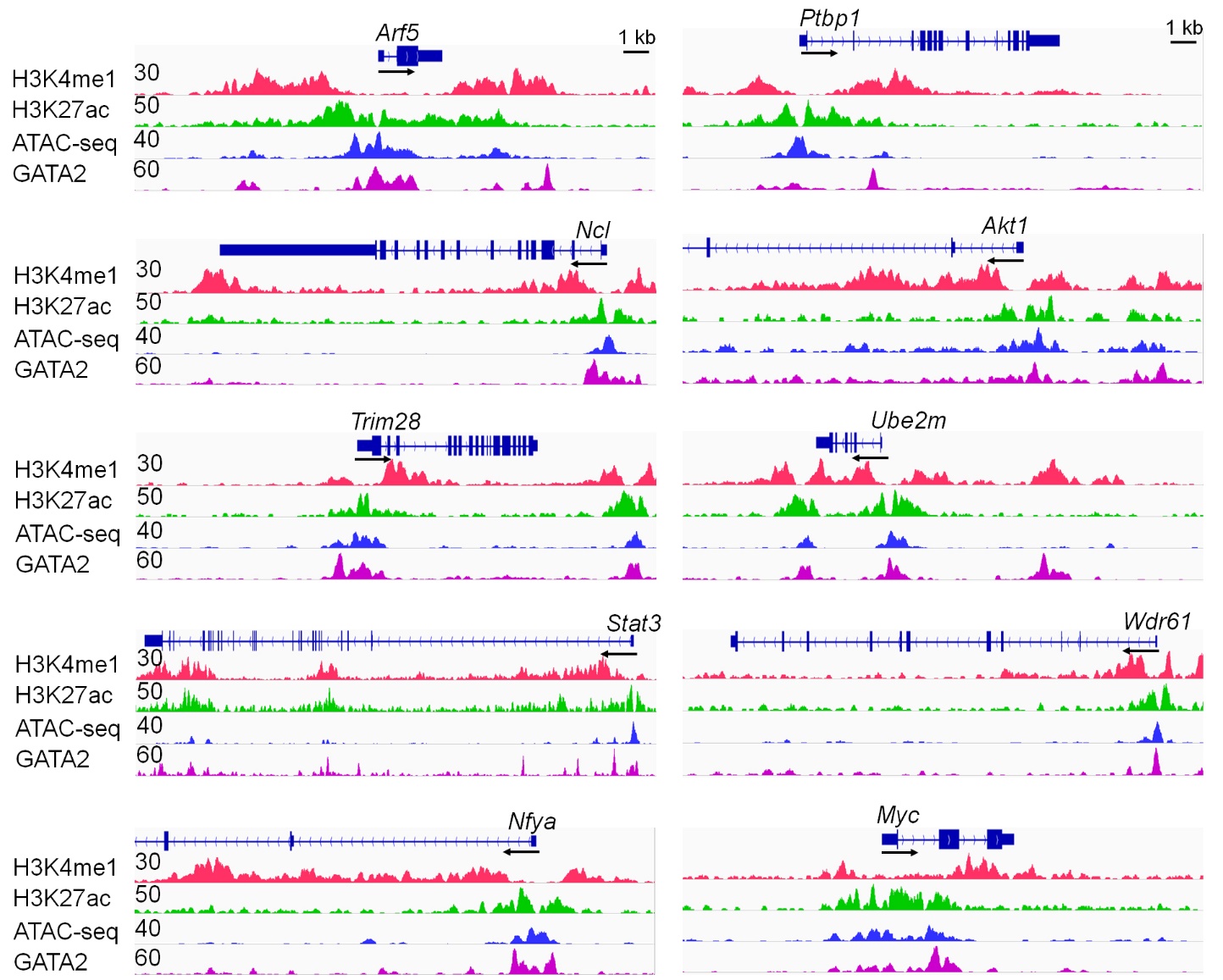

**Supplementary Figure 5 (related to Fig 3). GATA2 binding to the enhancers of genes identified as self-renewal genes in human embryonic stem cells and macrophages.** Representative H3K4me1, H3K27ac and GATA2 ChIP-seq tracks and Omni-ATAC-seq IGV tracks for the identified self-renewal genes. The representative ChIP-seq and ATAC-seq IGV tracks shown in the figure are generated from biological replicate 1 with similar patterns from biological replicate 2.

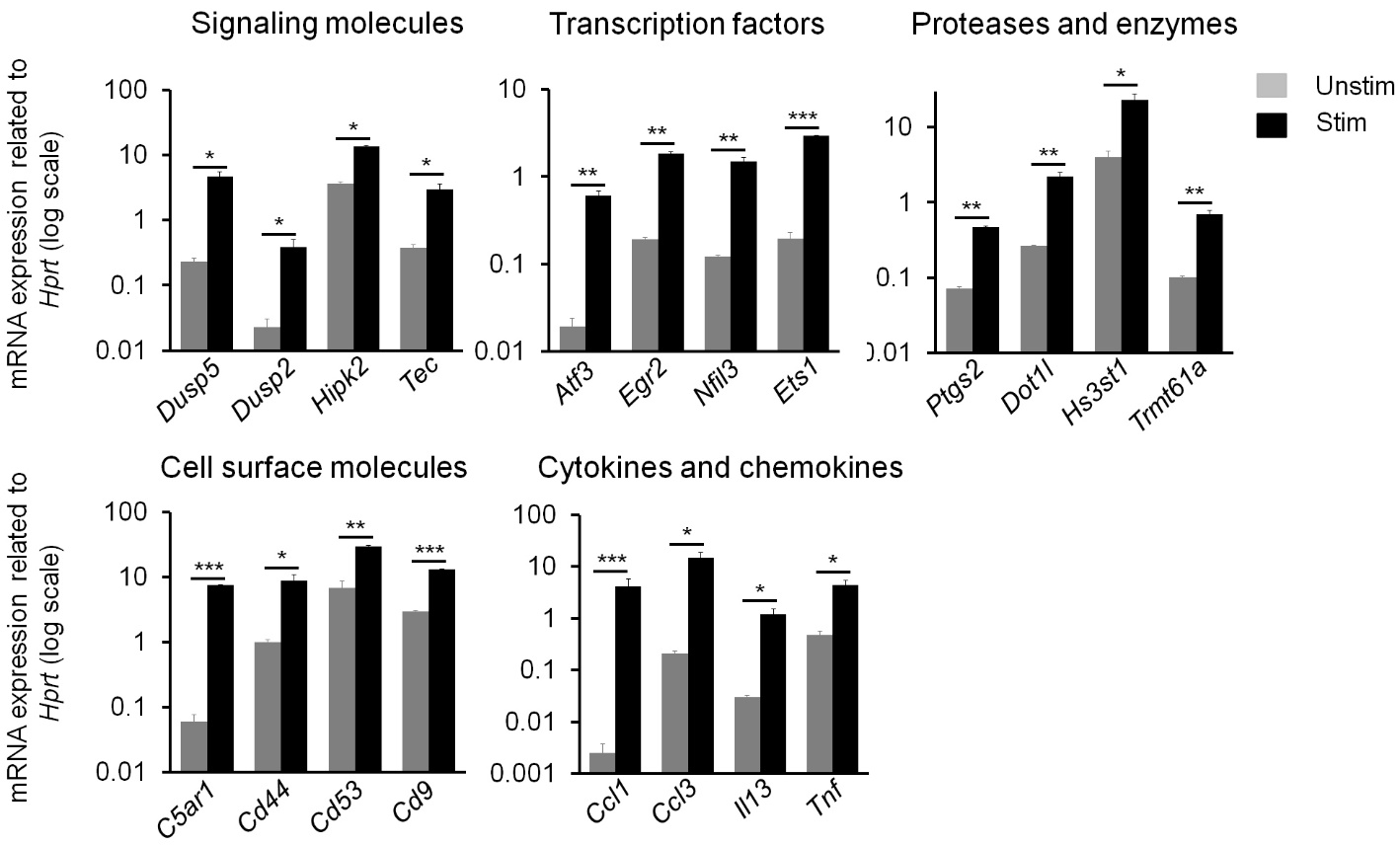

**Supplementary Figure 6 (related to Fig 4). qPCR validation of the expression of induced genes identified by RNA-seq data.** BMMCs were untreated (Unstim) or treated by IgE receptor crosslinking for two hours (Stim). qPCR analysis of the mRNA expression of activation-induced genes identified by RNA-seq. The activation-induced genes enriched in signaling molecules, transcription factors, proteases and enzymes, cell surface molecules, and cytokines and chemokines are shown. P values were calculated using the two-tailed student’s *t* test. Data represent three biological samples. Error bars represent mean ± SD. *: p<0.05; **: p<0.01; ***: p<0.001.

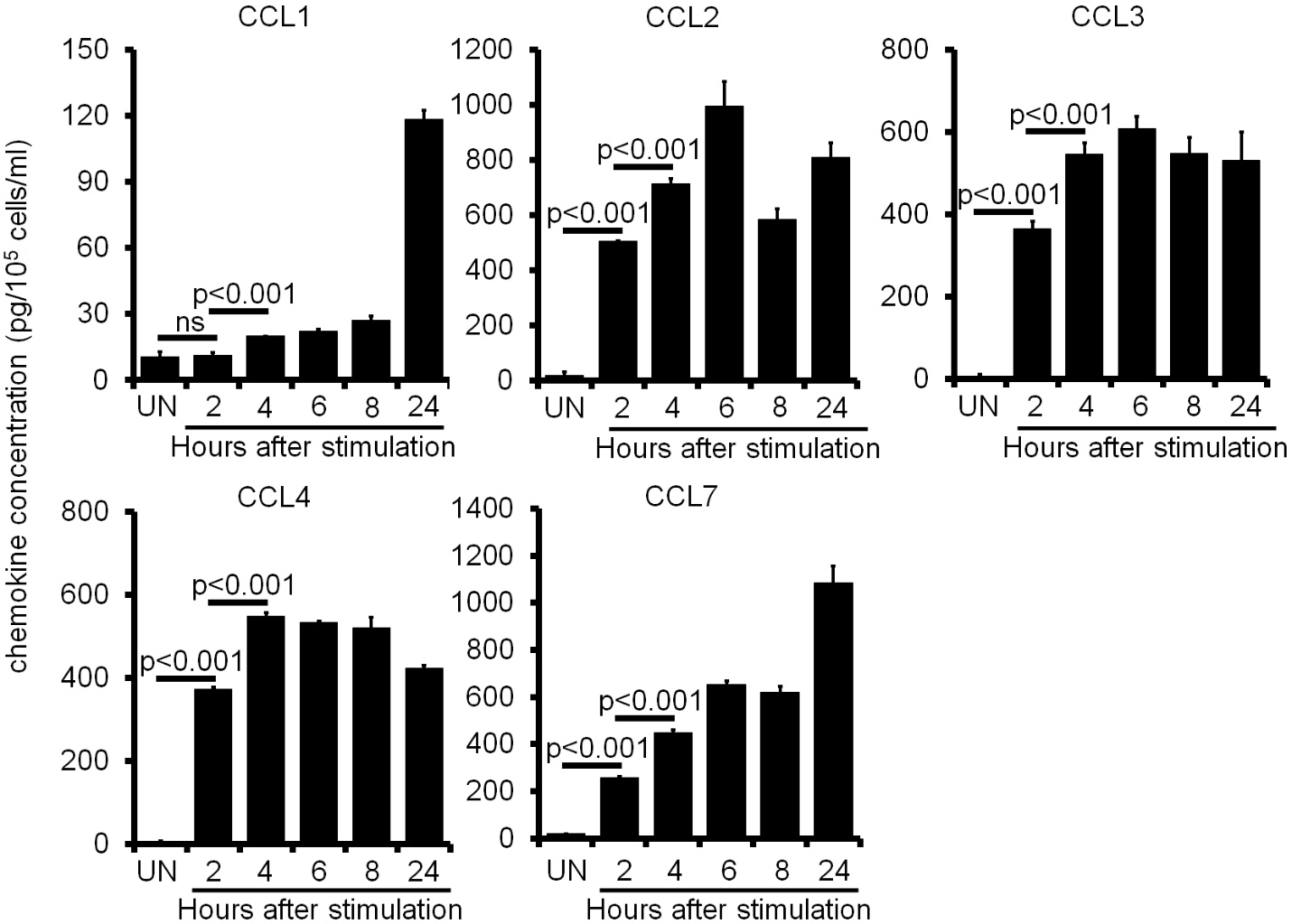

**Supplementary Figure 7 (related to Fig 4). ELISA analysis of the chemokine levels in resting and stimulated BMMCs.** BMMCs were untreated (UN) or treated by IgE receptor crosslinking for 2, 4, 6, 8 and 24 hours. Supernatants were collected and chemokine levels were analyzed by ELISA. P values were calculated using the two-tailed student’s *t* test. The data represents three biological samples. Error bars represent mean ± SD.

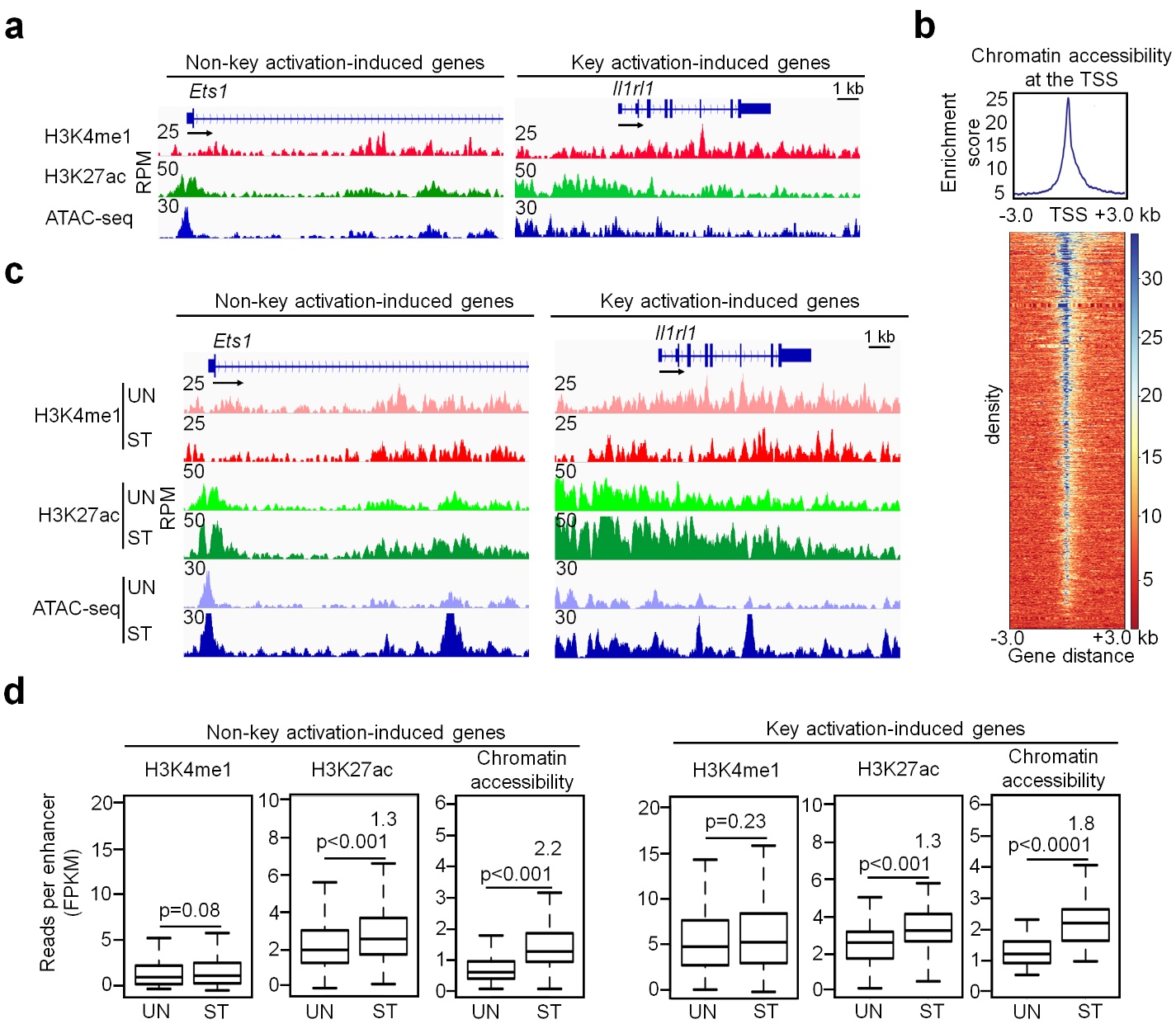

**Supplementary Figure 8 (related to Fig 6).**  **The activation-induced genes show “open” chromatin configuration before mast cells encounter antigen and show increased chromatin accessibility and H3K27ac modification in response to IgE receptor crosslinking. a** Representative IGV tracks of H3K4me1 ChIP-seq, H3K27ac ChIP-seq and ATAC-seq in the *Ets1* and *Il1rl1* gene loci in resting mast cells. **b** Enrichment distribution analysis of chromatin accessibility at the Transcription Start Sites (TSS) of the activation-induced genes. Chromatin accessibility enrichment scores were calculated for the regions at TSS ± 3kb of the cytokine and chemokine genes in resting mast cells using deepTools (v 3.3.0). **c** Representative IGV tracks of H3K4me1 ChIP-seq, H3K27ac ChIP-seq and ATAC-seq in the *Ets1* and *Il1rl1* gene loci in resting (UN) and activated (ST) BMMCs that were treated with IgE receptor crosslinking for two hours. **d** H3K4me1, H3K27ac and chromatin accessibility reads for the typical enhancers of non-key activation-induced genes and super-enhancers of key activation-induced genes in resting and activated mast cells. Reads were the sum of sequencing reads for all enhancers per gene and were normalized against enhancer sizes of the enhancers as FPKM. The numbers indicate the fold of induction of enhancer reads after IgE receptor crosslinking. P values were calculated using the two-tailed student’s *t* test. Data (**a**-**d**) represent two biological samples.

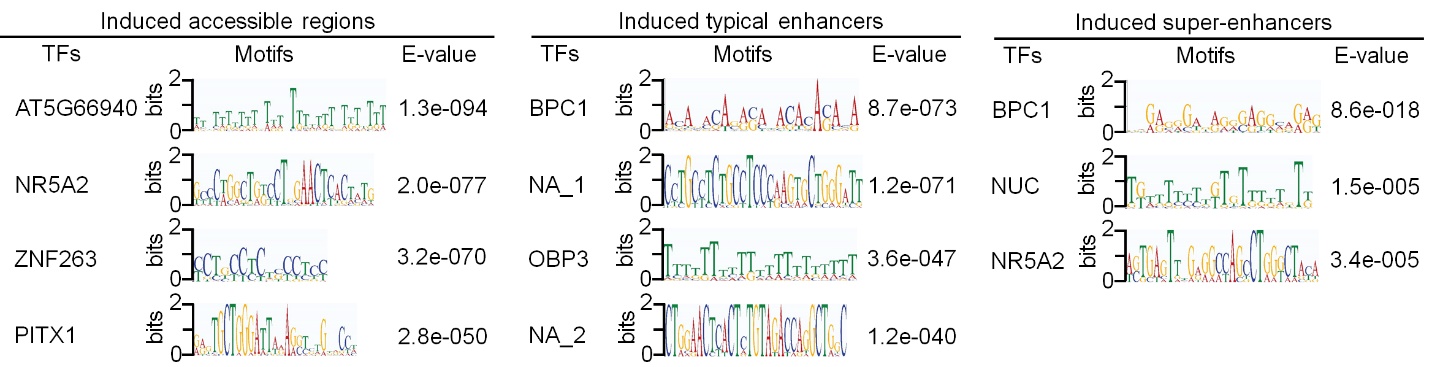

**Supplementary Figure 9 (related to Fig 7). Enrichment analysis of TF binding motifs.** TF binding motif enrichment analysis performed on the activation-induced accessible regions, activation-induced typical enhancers or activation-induced super-enhancers alone. NA means unknown TF binding motif.

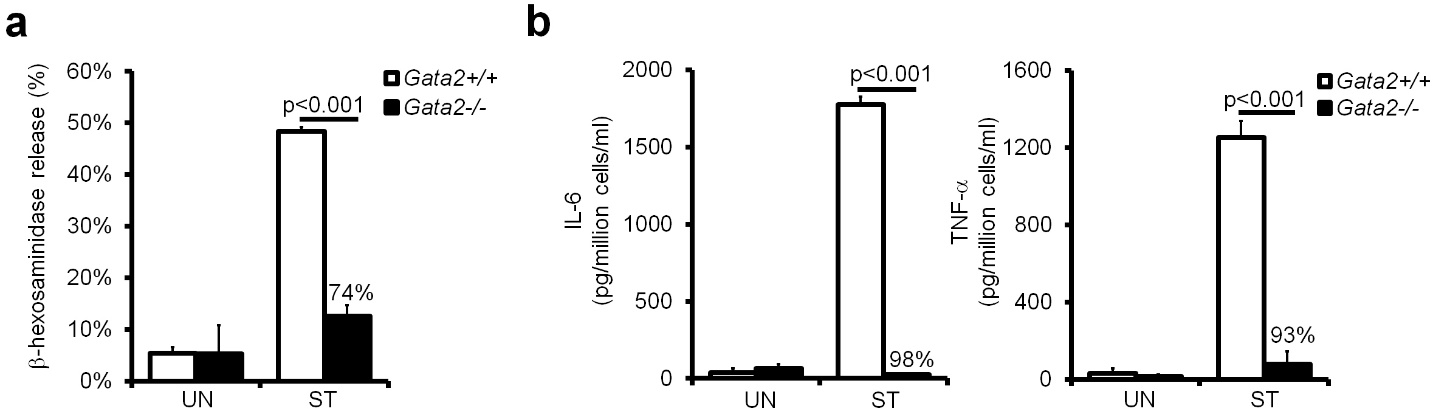

**Supplementary Figure 10 (related to Fig 7). ELISA analysis of the β-hexosaminidase release and IL-6 and TNF-α levels in resting (UN) and stimulated (ST) *Gata2* WT or KO BMMCs. a** *Gata2* WT or KO BMMCs were sensitized with 1 μg/mL IgE anti-TNP antibody for 24 hours and challenged with 100 ng/mL TNP-BSA for 30 minutes. Both cells and supernatants were collected. The percentage of β-hexosaminidase release was calculated as the percentage of β-hexosaminidase in supernatants relative to total β-hexosaminidase in supernatants and cell pellets. **b** *Gata2* WT or KO BMMCs were sensitized with 1 μg/mL IgE anti-TNP antibody for 24 hours and challenged with 100 ng/mL TNP-BSA overnight. The amounts of IL-6 and TNF-α protein in the supernatants were measured by ELISA. P values were calculated using the two-tailed student’s *t* test. Data represent three biological samples. The percentage indicates the reduction of β-hexosaminidase release and cytokine levels in *Gata2* WT BMMCs compared with *Gata2* KO BMMCs. Error bars represent mean ± SD.
